# Rich resource environment of fish farms facilitates phenotypic variation and virulence in an opportunistic fish pathogen

**DOI:** 10.1101/2020.06.10.144535

**Authors:** Katja Pulkkinen, Tarmo Ketola, Jouni Laakso, Johanna Mappes, Lotta-Riina Sundberg

**Author notes:** Corresponding author: Katja Pulkkinen, Address: Department of Biological and Environmental Science, P.O. Box 35, FI-40014, University of Jyväskylä, Finland.

## Abstract

Phenotypic variation allows adaptation of opportunistic pathogens to variable conditions in the outside-host environment with strong effects on their epidemiology and pathogenicity in hosts. Here we found that the isolates of an opportunistic fish pathogen *Flavobacterium columnare* from fish farming environment had higher phenotypic variation between two morphotypes in growth, as compared to the isolates from the natural water environment. The rough morphotypes had higher growth rate than the rhizoid morphotypes especially in the higher resource concentrations and in the higher temperature, but only if the isolate was originating from the fish farms. Rhizoid morphotype was more virulent than the rough type regardless of their origin. However, the virulence of the rough type increased sharply with the size of the fish, and the bacterial isolates from the gills of diseased fish were rhizoid type, indicating a reversal of the rough morphotype into rhizoid in contact with the fish. The high growth rate of the rough morphotype combined with the morphotype reversibility could increase the probability of columnaris epidemics at fish farms. Our findings suggest that intensive farming imposes different selection pressures on bacterial survival in the outside-host environment and its transmission compared to the natural water environment.

## Introduction

Phenotypic variation is one of the mechanisms that allows organisms to adapt to alternating environments (Via et al., 1995; Rainey and Travisano, 1998; Ackermann, 2015; D’Souza, 2020). This ability might be especially important for opportunistic pathogens that survive and replicate not only within the host but also in the outside-host environment (Schreiber et al., 2016). The ecological opportunities available outside the host generally differ vastly from those encountered by the pathogen inside the host (Brown et al., 2012; Anttila et al., 2016). Within the host, the greatest challenges are posed by the host immune system (Schmid-Hempel, 2009). In the outside host environment, low availability of resources, presence of predators, parasites and competitors, and abiotic factors, such as temperature, often restrict the growth (Friman et al., 2009; Adiba et al., 2010; Hibbing et al., 2010; Friman et al., 2011; Zhang et al., 2014). The ability to change from one alternative phenotype to another with different characteristics e.g. for competitive ability or immune evasion might be crucial for the expression of pathogenicity for opportunistic pathogens (Holland et al., 2014; Kreibich and Hardt, 2015; Ketola et al., 2016). In bacteria, phenotypic variation is often visible in the form of different types of colony morphologies, with differences in growth characteristics (Rainey and Travisano, 1998; Koh et al., 2007; Friman et al., 2009; Kunttu et al., 2009a; Sundell et al., 2013). Phenotypic variation in bacteria may rise as a response to change in environmental conditions (phenotypic plasticity) or as a consequence of stochastic switching between alternative phenotypes produced by the same genotype (phenotypic heterogeneity) (Ackermann, 2015; D’Souza, 2020).

Phenotypic variation might not, however, be only advantageous, but costs associated e.g. with maintaining the metabolic machinery for each environment, or trade-offs between different phenotypic properties in different environments may limit the adaptive value of phenotypic variation (Dewitt et al., 1998). However, these trade-offs may be relaxed if resource rich environment enables stronger allocation to growth and defense simultaneously (Friman et al., 2008). In addition, predictability and the speed of the environmental change determine the profitability of phenotypic change (Levins, 1969; Padilla and Adolph, 1996; DeWitt, 1998; DeWitt and Scheiner, 2004; Kussell and Leibler, 2005; Kristensen et al., 2008).

Intensive farming has been associated with increases in virulence for pathogens such as Marek’s disease virus in poultry farming (Atkins et al., 2013) as well as salmon lice (Mennerat et al., 2010; Mennerat et al., 2012; Ugelvik et al., 2017) and *Flavobacterium columnare* in aquaculture (Pulkkinen et al., 2010; Sundberg et al., 2016). The conditions and practices in intensive farming are likely to select for different genotypic and/or phenotypic properties than the conditions outside these environments. For pathogens, intensive farming represents environment with more possibilities and challenges for growth and survival. On one hand, high density of genetically homogeneous hosts and excess feed offer abundant resources for growth, but on the other, e.g. medical treatments during outbreak season pose a recurrent and strong threat for pathogen survival (Mennerat et al., 2010). In pathogens, such environment could select increased phenotypic variation for coping in the both extremes of the outside-host environment.

In this work, we studied phenotypic variation in an opportunistic fish pathogen *Flavobacterium columnare* isolated either from fish farms or from natural waters in Finland. The bacterium is the causative agent of columnaris disease, which causes severe economic losses in freshwater fish farming affecting e.g. channel catfish (*Ictalurus punctatus*) in US (Wagner et al., 2002) and salmonid production in Europe (Declercq et al., 2013). It has been suggested that the virulence of the bacterium has increased during the last few decades at fish farms (Pulkkinen et al., 2010; Sundberg et al., 2016; Ashrafi et al., 2018). Growth and replication in the outside-host environment (Kunttu et al., 2009b; Sundberg et al., 2014; Pulkkinen et al., 2018) exposes the bacterium to selection pressures not driven by the host-pathogen interaction. For example, in the farming environment the use of chemical and antibiotic treatments makes the environment more unpredictable than the natural conditions, and can directly influence bacterial growth. The pontaneous change of colony types expressed by *F. columnare* is reversible. While the exact mechanisms leading to colony morphology changes are not known, the different colony types are suggested to serve different functions in invasion and replication in fish and in the outside host environment (Kunttu et al., 2009a; Kunttu et al., 2009b; Laanto et al., 2014; Zhang et al., 2014).

We hypothesized that in order to maximize survival, isolates of fish farm origin would present higher phenotypic variation than isolates from natural waters and that these differences would be more pronounced in higher resources and higher temperature. In order to test the hypothesis, we measured the growth of two morphotypes (ancestral rhizoid, Rz, and its rough, R, derivative, Figure S1) of each of five isolates originating from both environments in five different resource concentrations and in two temperatures, one corresponding to optimal growth temperature and the other below the optimum (Ashrafi et al., 2018). We also tested the stability of the morphotypes in plate cultivation under the resource concentrations used for testing the growth. Finally, we hypothesized that the bacteria isolated from fish farms would have higher virulence, which we tested in fish challenge tests *in vivo*.

## Results

### Growth rate

Maximal bacterial growth rate was found to be affected by several factors (Tables 1 and 2). Growth rate was affected the most by the temperature, with higher temperature supporting higher growth rate (Table 2). Growth rate also increased with resource concentration (0.05x, 0.1x, 0.5x, 1x and 2x growth medium), although there were no statistical differences between the two lowest resource concentrations (p<0.3 in pairwise tests). The rest of the pairwise tests indicated differences between resource concentrations (p<0.001). The origin of isolate or morphotype affected growth rate only in interaction with other factors. Within fish farm isolates, rough morphotype (R) replicated faster than the rhizoid (Rz) (Figure 1a, F_1,620_=10.048, p=0.002), whereas this was not evident within isolates from natural water (F_1,620_=1.902, p=0.168), supporting our hypothesis of higher phenotypic variation in the fish farm isolates. Moreover, growth rate of fish farm isolates was higher than the growth rate of isolates from natural waters only in the R morphotype (F_1,620_=9.772, p=0.010), but not in the Rz morphotype (F_1,620_=0,620, p=0.448). Growth advantage of R morphotype was found only at high temperature (F_1,772_=7.551, p=0.006, Figure 1b) whereas at low temperature there were no differences between morphotypes (F_1,772_=0.916, p=0.339).

**Table 1.**
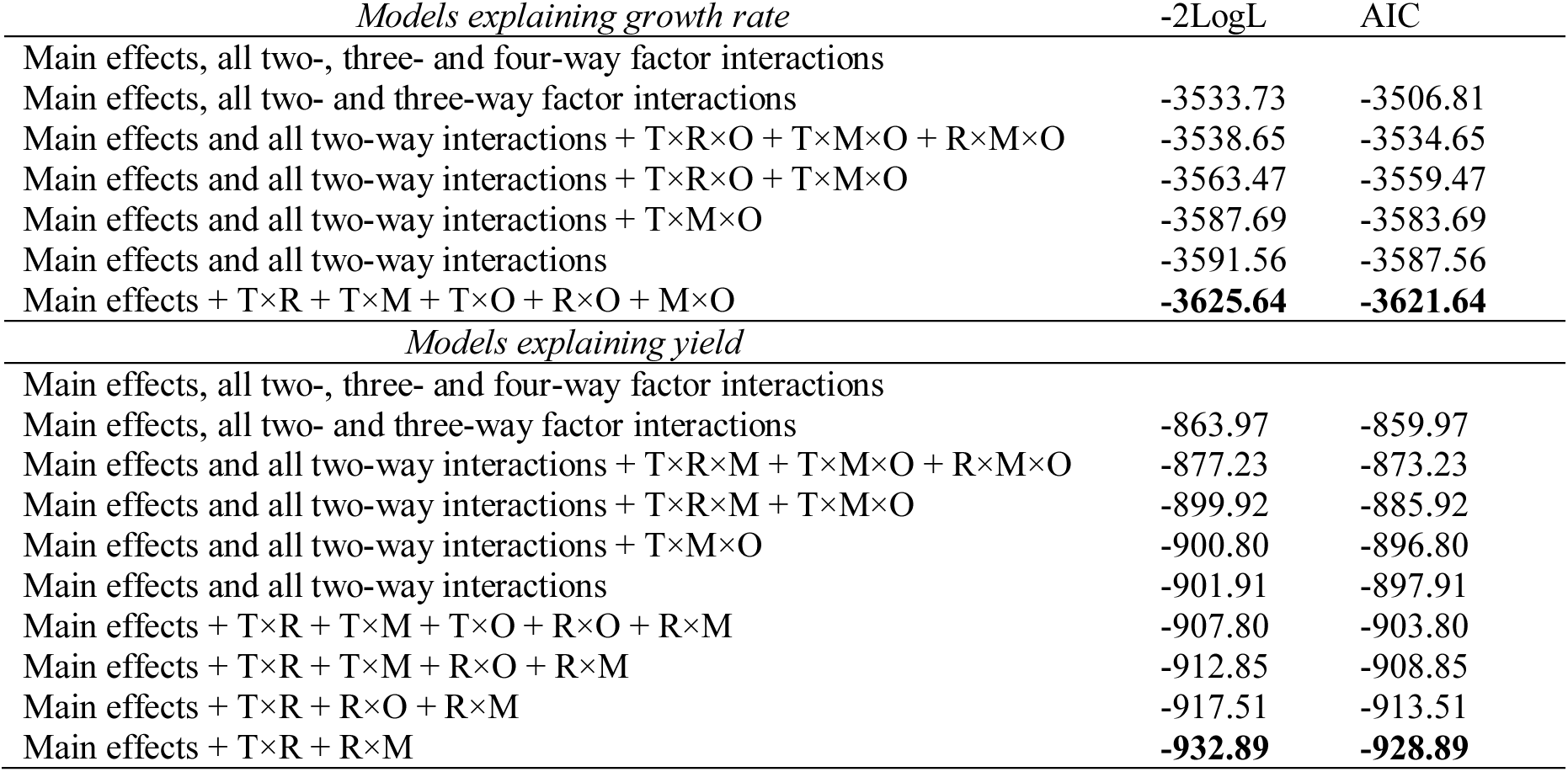
Model selection for models explaining growth rate and yield based on AIC-criteria of fitted models (smaller the value the better the model). Model fitting was done with backward model reduction by excluding non-significant parameters one by one, starting from highest order interactions. Models included main effects of temperature (T), resource concentration (R), morphotype (M) and origin of isolate (O), and their interactions. All models contained also fixed effect of measurement block (measurement week) and random effect of isolate’s identity, to control for non-independency of observations. Best models are highlighted with bold.

| *Models explaining growth rate* | -2LogL | AIC |
| --- | --- | --- |
| Main effects, all two-, three- and four-way factor interactions |  |  |
| Main effects, all two- and three-way factor interactions | -3533.73 | -3506.81 |
| Main effects and all two-way interactions + T×R×O + T×M×O + R×M×O | -3538.65 | -3534.65 |
| Main effects and all two-way interactions + T×R×O + T×M×O | -3563.47 | -3559.47 |
| Main effects and all two-way interactions + T×M×O | -3587.69 | -3583.69 |
| Main effects and all two-way interactions | -3591.56 | -3587.56 |
| Main effects + T×R + T×M + T×O + R×O + M×O | **-3625.64** | **-3621.64** |
| *Models explaining yield* |  |  |
| Main effects, all two-, three- and four-way factor interactions |  |  |
| Main effects, all two- and three-way factor interactions | -863.97 | -859.97 |
| Main effects and all two-way interactions + T×R×M + T×M×O + R×M×O | -877.23 | -873.23 |
| Main effects and all two-way interactions + T×R×M + T×M×O | -899.92 | -885.92 |
| Main effects and all two-way interactions + T×M×O | -900.80 | -896.80 |
| Main effects and all two-way interactions | -901.91 | -897.91 |
| Main effects + T×R + T×M + T×O + R×O + R×M | -907.80 | -903.80 |
| Main effects + T×R + T×M + R×O + R×M | -912.85 | -908.85 |
| Main effects + T×R + R×O + R×M | -917.51 | -913.51 |
| Main effects + T×R + R×M | **-932.89** | **-928.89** |

**Table 2.** Results of mixed model analyses on determinants of growth rate and yield in *Flavobacterium columnare* isolates. Clones were either isolated from fish farms or from natural waters. From each clone we measured two morphotypes (rough and rhizoid) in high and low temperatures (25 °C and 15 °C) and in five different resource concentrations (0.05x, 0.1x, 0.5x, 1x and 2x Shieh-medium). In statistical analyses we also included effects of measurement block (identical measurements were done in two subsequent weeks), and maternal isolate’s identity to control for non-independence of observations arising from shared genetic background. Excluded factor interactions are denoted with -.

|  | *Growth Rate* |  |  |  | *Yield* |  |  |  |
| --- | --- | --- | --- | --- | --- | --- | --- | --- |
|  | df1 | df2 | F | p. | df1 | df2 | F | p. |
| Temperature | 1 | 772 | 1298.417 | <0.001 | 1 | 775 | 709.919 | <0.001 |
| Resource Concentration | 4 | 772 | 168.453 | <0.001 | 4 | 775 | 1527.139 | <0.001 |
| Morphotype | 1 | 772 | 4.411 | 0.069 | 1 | 775 | 6.621 | 0.010 |
| Origin of Isolate | 1 | 8 | 1.603 | 0.206 | 1 | 8 | 0.017 | 0.899 |
| Temperature × Resource | 4 | 772 | 62.830 | <0.001 | 4 | 775 | 74.572 | <0.001 |
| Temperature × Morphotype | 1 | 772 | 6.863 | 0.009 | - | - | - | - |
| Temperature × Origin of Isolate | 1 | 772 | 17.886 | <0.001 | - | - | - | - |
| Resource × Morphotype | - | - | - | - | 4 | 775 | 6.984 | <0.001 |
| Resource × Origin of Isolate | 4 | 772 | 2.788 | 0.026 | - | - | - | - |
| Morphotype × Origin of Isolate | 1 | 772 | 10.346 | <0.001 | - | - | - | - |
| Block | 1 | 772 | 34.581 | <0.001 | 1 | 775 | 2.958 | 0.086 |
| Isolate's ID | $\sigma^2=3.43 \times 10^{-5}$, Wald Z=1.695, p=0.090 | | | | $\sigma^2=5.47 \times 10^{-3}$, Wald Z=1.931, p=0.053 | | | |

**Figure 1.**
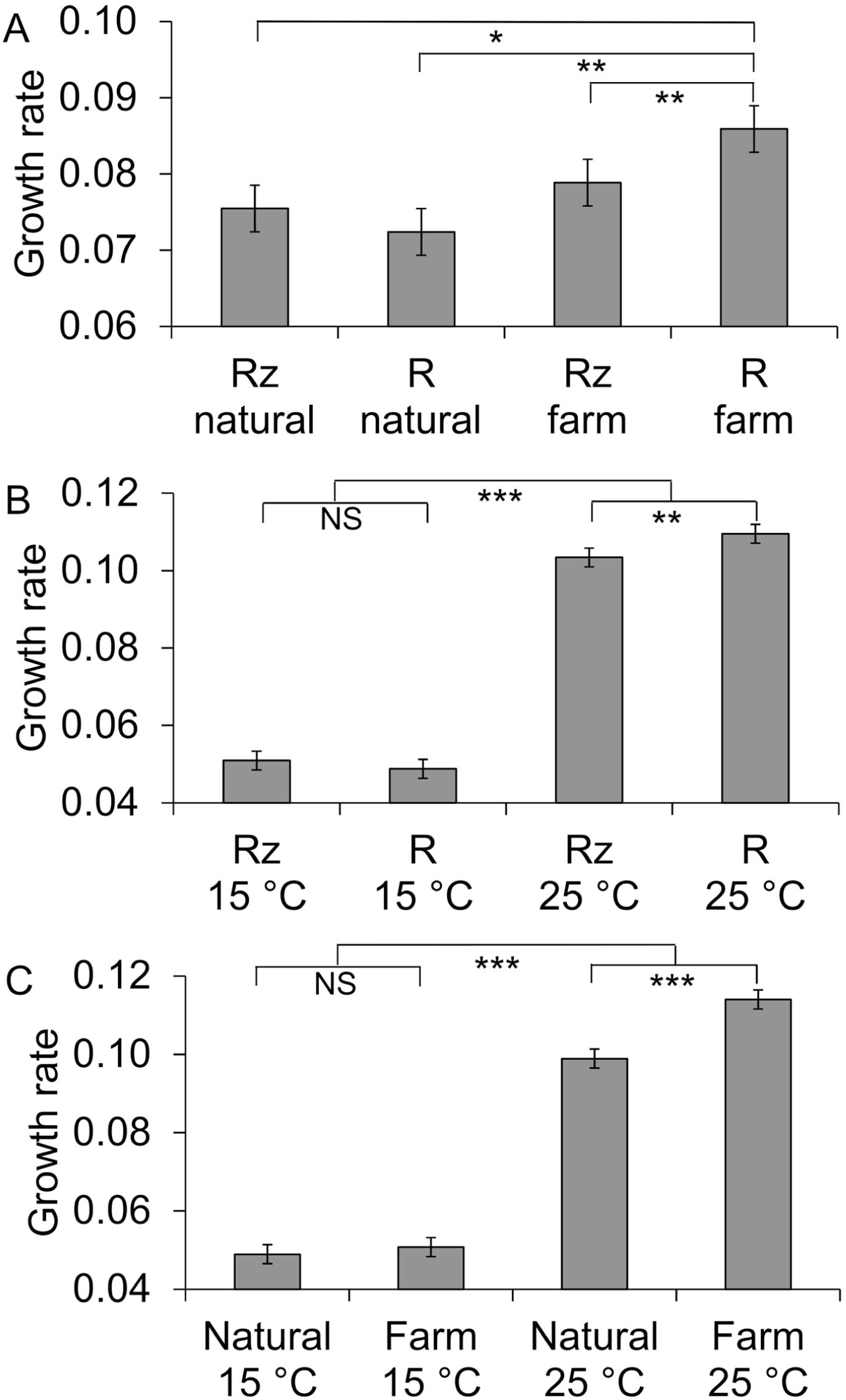
Effects of morphotype (Rz = rhizoid, R = rough) and origin (Natural = natural waters, Farm = fish farm) (A), morphotype and temperature (B) and origin of isolation and temperature (C) on maximal growth rate of *Flavobacterium columnare*. Significant differences between treatments are denoted with *, **, ***; p<0.05, p<0.01, p<0.001, respectively.

Farm isolates had higher growth rates than isolates from natural water in high temperature (Figure 1c, F_1,10.620_=12.209, p=0.005), but not in low temperature (F_1,10.620_=0.176, p=0.684). Farm isolates had significantly higher growth rates than isolates from natural waters in the two highest resource concentrations (Figure 2a, 0.05x medium: F_1,20.637_=0.826, p=0.374; 0.1x: F_1,20.637_=0.906, p=0.352; 0.5x: F_1,20.637_=0.630, p=0.436; 1x: F_1,20.637_=11.882, p=0.002, 2x: F_1,20.637_=4.740, p=0.041).

**Figure 2.**
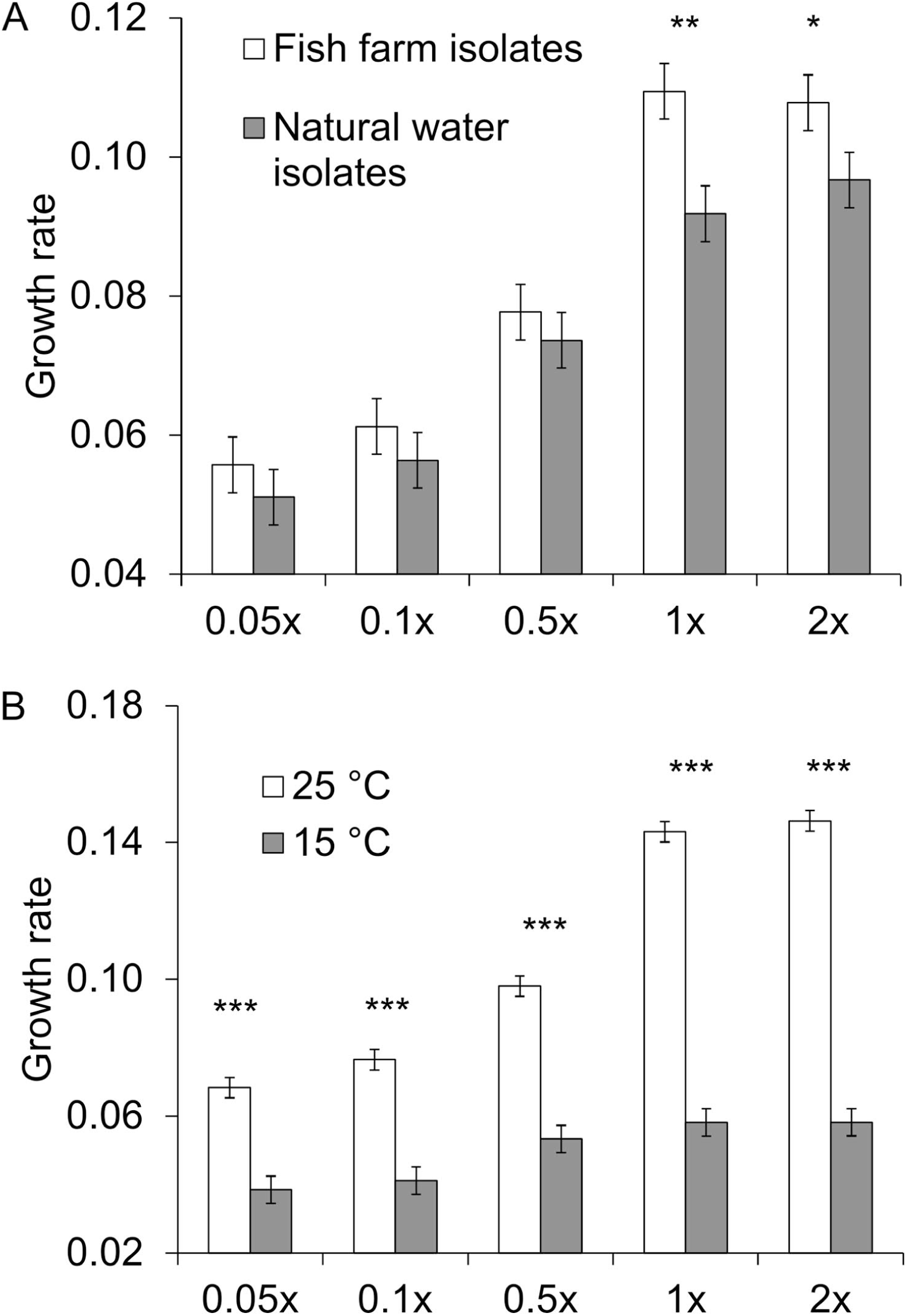
Effects of resource concentration (0.05x, 0.1x, 0.5x, 1x and 2x Shieh medium) and origin of isolate (A) and temperature (B) on maximal growth rate of *Flavobacterium columnare*. Significant differences between treatments are denoted with *, **, and ***; p<0.05, p<0.01, and p<0.001, respectively).

Differences in growth rates between resource concentrations were larger in high temperature (Figure 2b): in high temperature all pairwise tests between different resource concentrations were clearly significant (p<0.006) except 0.05x vs 0.1x (p=0.203) and 1x vs. 2x (p=0.999). In low temperature differences between 0.05x and 0.1x (p=0.999), 0.5x vs. 1x (p=0.999), 0.5x vs. 2x (p=0.999) were not found, whereas other pairwise comparisons yielded significant differences.

### Yield

We found that biomass yield was best explained by a model with significant main effects of temperature, colony morphology and medium, and interactions between medium and temperature, and medium and colony morphology (Table 1, 2). Higher temperature led to higher biomass yield than lower temperature (yield at 15 °C= 0.492, s.e.=0.024; 25 °C= 0.728, s.e.=0.024, Table 2). In addition, Rz colony morphology (yield=0.621, s.e.=0.024) had higher biomass yield than R colony morphology (yield=0.599, s.e.=0.024, Table 2). The biomass yield was higher the richer the medium (p<0.001 for all pairwise tests between different resource concentrations).

Rz morphotypes produced the highest biomass yield in intermediate resource concentrations (0.5x: F_1,775_=8.795, p=0.003; 1x: F_1,775_=22.882, p<0.001, Figure 3a). In the smallest and in the largest concentrations, colony types did not differ in their biomass yield (0.05x: F_1,775_=0.105, p=0.746, 0.1x: F_1,775_=2.773, p=0.096, 2x: F_1,775_=0.001, p=0.995). Increase in resource concentration increased biomass yield, and the increase was larger in high temperatures (Figure 3b, Table 2. 0.05x resource: F_1,775_=0.688, p=0.407, 0.1x: F_1,775_=22.458, p<0.001, 0.5x: F_1,775_=273.742, p<0.001, 1x: F_1,775_=437.327, p<0.001, 2x: F_1,775_=273.993, p<0.001).

**Figure 3.**
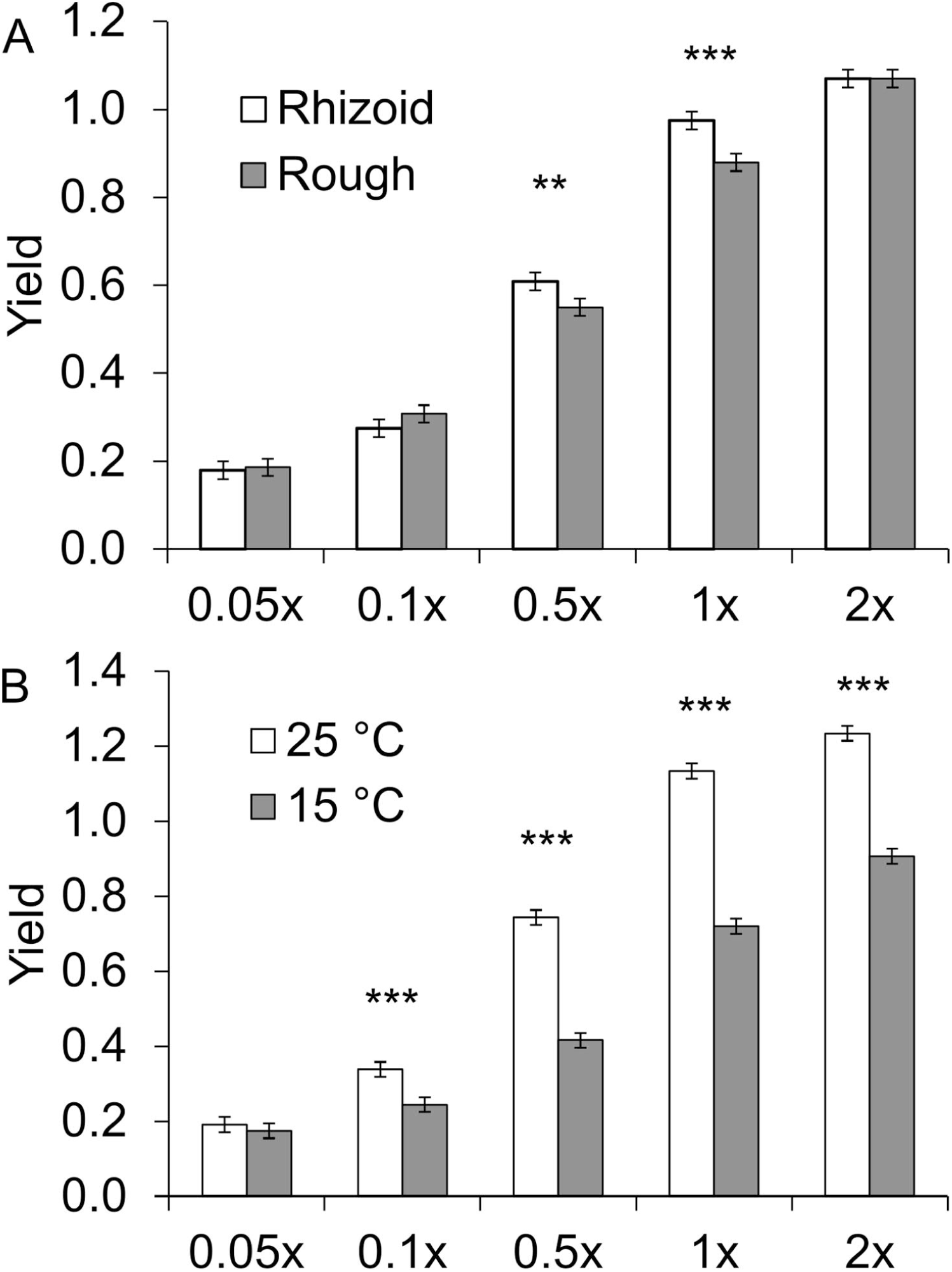
Effects of resource concentration (0.05x, 0.1x, 0.5x, 1x and 2x Shieh medium) and morphotype (A), and resource concentration and temperature (B) on maximal biomass yield of *Flavobacterium columnare*. Significant differences between treatments are denoted with *, **, ***; p<0.05, p<0.01, p<0.001, respectively.

### Morphotype stability

When plated on agar plates with the four highest (0.1x, 0.5x, 1x, 2x) nutrient concentrations used in the growth experiment, some of the originally rough colony morphotypes grew in a rhizoid form in the concentrations from 0.1x to 1x. At the highest resource concentration (2x), the colony spreading decreased, making also the rhizoid colonies resemble rough ones (Figure S2). This change was more pronounced for isolates from natural waters (Fisher’s exact test, p = 0.043) than for the fish farm isolates (p = 0.090; Figure 4). Plating on 0.05 x Shieh agar did not result in visible colonies for most bacterial cultures, hence this treatment was omitted from statistical analysis.

**Figure 4.**
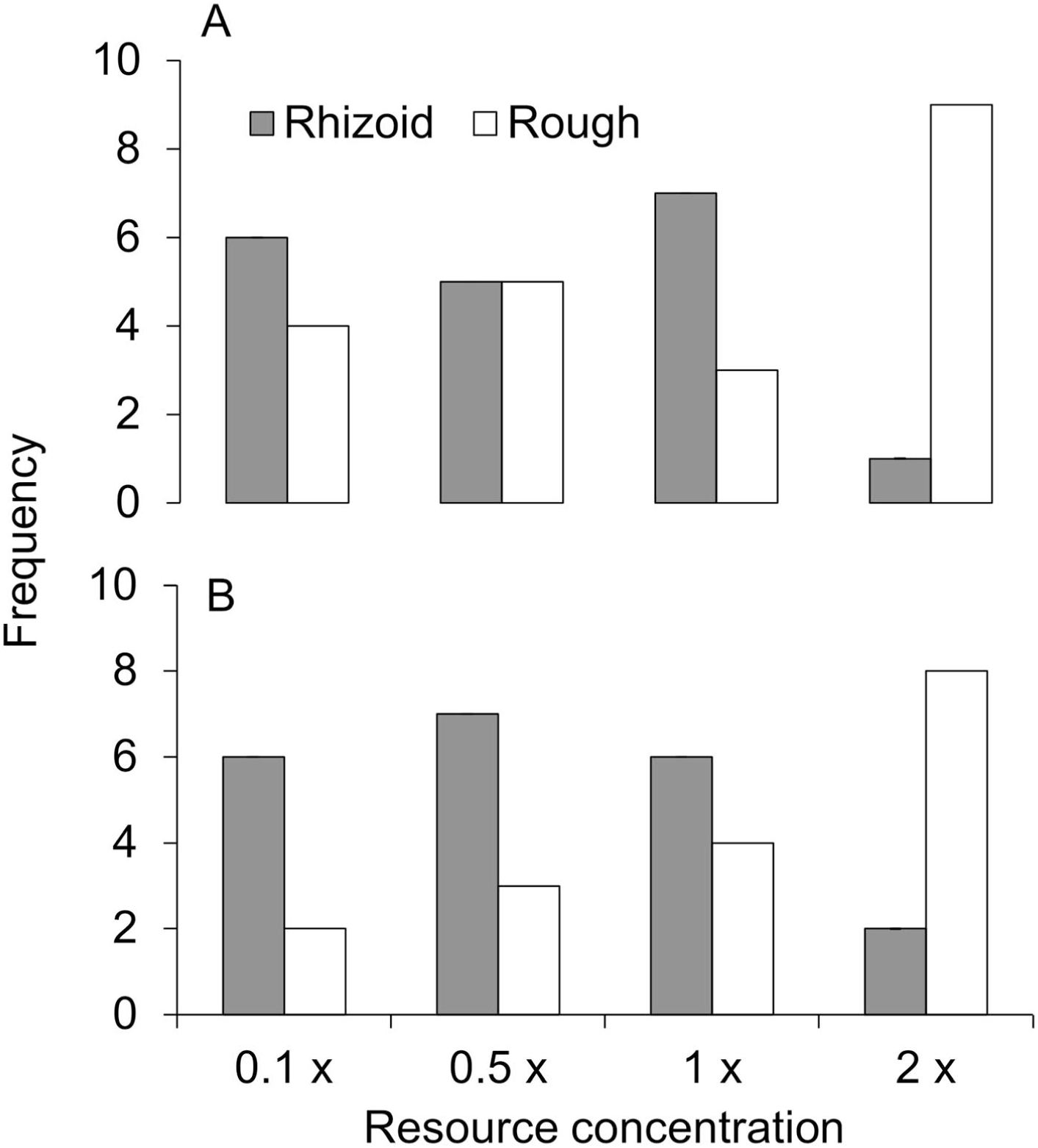
Stability of the rhizoid and rough colony morphotype on agar plates containing different concentrations of Shieh growth medium (0.1x, 0.5x, 1x and 2x) for five isolates from natural waters (A) and five fish farm isolates (B). Note that one fish farm isolate did not grow on 0.1 x Shieh concentration.

### Virulence test

In contrast to our hypothesis, isolate origin did not affect virulence. The best model explaining fish morbidity in the challenge experiment included the effects of colony morphology, fish weight and their interaction (Table 3). The virulence was higher for the rhizoid morphotype than for the rough morphotype and increased with fish weight. For rhizoid colony type, the morbidity risk increased steadily with fish size, while the morbidity risk induced by the rough colony morphology remained lower than that induced by the rhizoid type until a sharp rise at the largest fish sizes (Table 4, Figure 5). The isolates from gills of diseased fish were always rhizoid, also for those fish that were exposed to rough morphotype. All control fish survived until the end of the experiment. No *F. columnare* could be isolated at the end of the experiment from those fish who did not exhibit disease symptoms or from control fish. Rough morphotypes of the isolates B392 and B067 did not cause disease in exposed fish.

**Table 3.** Model selection of the virulence experiment based on Akaike information criteria (AIC). The best fit model estimating morbidity of the host (zebra fish) within time is marked with bold. O: origin (nature, fish farm), T: type (rhizoid, rough), W: fish weight (g), S: strain identity, + indicates the main effects,: indicates interaction.

| Model | AIC | DF |
| --- | --- | --- |
| O+T+W+O:T+O:W+T:W+O:T:W+(1\|S) | 488.9 | 9 |
| O+T+W+O:T+O:W+T:W+(1\|S) | 487.41 | 8 |
| O+T+W+O:W+T:W+(1\|S) | 485.44 | 7 |
| O+T+W+O:T+T:W+(1\|S) | 485.41 | 7 |
| O+T+W+O:T+O:W+(1\|S) | 494.05 | 7 |
| O+T+W+T:W+(1\|S) | 483.44 | 6 |
| **T+W+T:W+(1\|S)** | **481.72** | **5** |
| T+W+(1\|S) | 488.26 | 4 |

**Table 4.** The effect of colony morphology (type) and fish weight on the morbidity risk of the host fish in the virulence experiment.

| Source | Estimate | SE | df | Chi sq. | p |
| --- | --- | --- | --- | --- | --- |
| Intercept | -3.05 | 0.3 |  |  |  |
| type (rough) | -2.1 | 0.41 | 1 | 37.67 | <0.001 |
| weight | 1.08 | 0.62 | 1 | 20.99 | <0.001 |
| type x weight | 3.04 | 1.02 | 1 | 8.83 | 0.003 |

**Figure 5.**
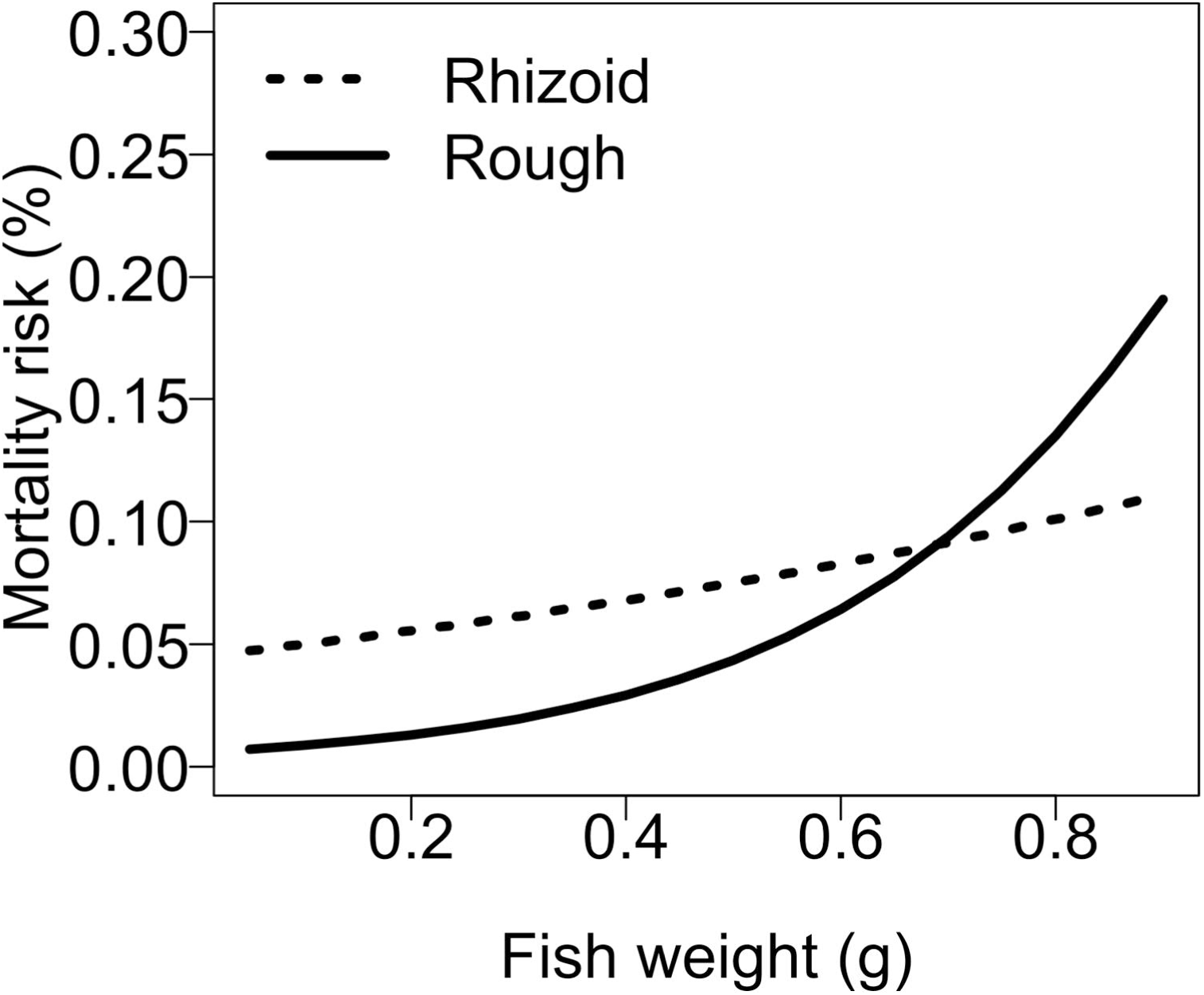
The estimated mortality risk per hour for zebra fish infected with rhizoid or rough morphotypes of *Flavobacterium columnare*.

## Discussion

Genetically identical microbes often show substantial variation in morphology and various life-history traits (Rainey and Travisano, 1998; Ackermann, 2015; D’Souza, 2020). It has been suggested that such variation accelerates the rate of adaptive evolution in populations facing novel environmental challenges. To understand how different environmental challenges have facilitated phenotypic variation of an opportunistic pathogen *F. columnare*, we compared the growth features and virulence of different morphotypes of strains originating from fish farms and from natural waters. We found that the rough morphotypes of fish farm isolates had higher growth rate than the rhizoid morphotypes originating from fish farm, or either of the morphotypes of isolates originating from natural waters (Figure 1A). Moreover, the higher growth rate of the rough morphotype, as compared to the rhizoid type, was pronounced in higher resource concentrations and in the higher temperature. These findings support our hypothesis that fish farms impose different selection pressures on bacterial survival and transmission in the outside-host environment than natural waters. In contrast to our hypothesis, we did not find difference in virulence based on the origin of the isolates, but the rhizoid type was more virulent than the rough type. Interestingly, the bacterial isolates from gills of diseased fish exposed to either morphotype exhibited only rhizoid morphotypes. Moreover, rough morphotypes of several isolates induced host death, suggesting that the rough morphotype reverted to rhizoid once in contact with the fish.

The bacterial growth rates were almost four times higher at 25°C, which falls within the optimum temperature for growth for *F. columnare* isolates in Finland (range 23.7-27.9°C) (Ashrafi et al., 2018), than below the optimum at 15°C. In addition, the yields increased with temperature. *F. columnare* is considered a warm water species worldwide (Declercq et al., 2013), and the disease outbreaks occur only during summer, when water temperatures reach c.a. 18°C at Finnish fish farms (Pulkkinen et al., 2010). The higher temperature increased the growth of the fish farm isolates more than the isolates from natural waters, which could be a yet another indication of improved ability of fish farm isolates to respond to optimal growth conditions. In addition, increase in resource concentration affected growth only in the higher temperature, highlighting the importance of temperature for fitness of *F. columnare*.

Increase in resource concentration improved both the growth rate and yield of the bacteria. For yield, the rhizoid morphotype attained higher values than the rough morphotype, but only at two intermediate resource concentrations (0.5x and 1x), with not effect of origin of the isolate. Growth rate and efficiency of resource utilization (yield) are opposing metabolic strategies that are thought to be traded-off with each other in microbes (Novak et al., 2006). However, the statistical analyses of the results of our experiment suggested that differences between treatments were not caused by life-history tradeoffs between maximal growth rate and yield.

For growth rate, the rough morphotype originating from fish farms had higher growth rates with increasing temperature and resource concentration than rough morphotypes of natural water isolates or rhizoid morphotypes of either origin. At fish farms, resource concentrations in the outside-host environment are high due to fish feeding and accumulation of feces. This is evident particularly in increased temperatures when the feeding activity of the fish is impaired, further elevating the concentration of uneaten fish feed and excreta in the water (Wedemeyer, 1996; Ellis et al., 2002). Thus, our findings could indicate that the rough morphotypes of fish farm isolates are adapted for utilizing increased resource concentrations in water at farms during warm water periods. Results pointing to this direction were found in a previous study, which found that *F. columnare* isolates in outlet water of a fish farm responded more to differences in resource concentration than isolates from inlet water (Sundberg et al., 2016).

Previous studies found that rhizoid morphotype is a prerequisite for *F. columnare* infectivity (Kunttu et al., 2009a; Laanto et al., 2012; Zhang et al., 2014). In our experiment, the mortality induced by the rough morphotype was clearly higher than in previous experiments testing differences between morphotypes (Kunttu et al., 2009a; Laanto et al., 2012; Laanto et al., 2014; Zhang et al., 2014), although still significantly lower than mortality induced by the rhizoid morphotype. The virulence in *F. columnare* increases with dose and increased resource concentrations in the outside-host environment (Penttinen et al., 2016; Kinnula et al., 2017). As all fish exposed to rough morphotype exhibited only rhizoid morphotypes in gill samples, our results suggest that rough morphotypes may play an important role for development of disease epidemics. By having a more rapid growth in high resource conditions, rough types could quickly give rise to a bacterial population expressing virulence in contact with the fish. Indeed, in rough morphotype, there was a tendency for positive correlation (Spearman’s rho = 0.7, n = 10, p =0.013, data not shown) between virulence and growth rate measured at 1x Shieh at 25°C (conditions used for fish challenge), suggesting that phenotypic change to virulent morphotype was connected to fast replication. For the rhizoid morphotype correlation between growth rate and virulence was not found. The virulence induced by rough morphotype increased sharply with the size of fish, whereas the virulence of the rhizoid morphotype increased more steadily with the fish size. This could also be an indication of association between growth rate and virulence, as larger fish hosts could either release more nutrients in the water or provide a larger surface area for bacterial attachment and growth.

The mechanisms of phenotypic variation for *F. columnare* are not known. Rough morphotype can appear spontaneously in serial cultivation (Kunttu et al., 2009a; Kunttu et al., 2012; Sundberg et al., 2014), while primary isolates from diseased fish from fish farms are always rhizoid type (Laanto et al., 2012). Rough morphotype is induced also as a response to phage infection (Laanto et al., 2012). The positive association between virulence and growth rate found between spontaneous rough morphotypes in this study could indicate phenotypic variation via random phenotype switch, as increase in number of replicative events (i.e. higher growth rate) facilitates a chance for such switch (Wick et al., 2002). In clonally replicating bacteria, the heterogeneity of the phenotype can be achieved by stochastic switching between alternative phenotypes produced by the same genotype (Balaban et al., 2004; Veening et al., 2008; Edwards, 2012; Ackermann, 2015; D’Souza, 2020). This strategy of bet-hedging buffers against the risks associated with inducible response to often unpredictable environmental clues (Kussell and Leibler, 2005; Schreiber et al., 2016).

The phenotypic variation for *F. columnare* has also been suggested to be regulated by gene expression as a plastic response to environmental change (Laanto et al., 2012; Penttinen et al., 2018). In our experiment, plating on agar containing the four highest resource concentrations used in the growth experiment, indicated instability in the rough morphotype, and variation between individual isolates in morphotype stability, confirming results of a previous study (Laanto et al., 2012). Similarly to previous studies (Laanto et al., 2012)(Penttinen et al., 2018), we also found that the spreading of the rhizoid morphotypes decreased in the highest resource concentration, resulting in colonies resembling the rough type, while some of the rough morphotypes reverted to rhizoid type in three of the lowest resource concentrations tested (Figure 4). Together these results suggest that morphotype change is inducible and related to nutrient scavencing, such that the rhizoid motile type is connected to finding scarce resources and the rough type adapted to fast utilization of rich resources. Further support comes from gene expression studies showing that genes associated with gliding motility increased expression under low nutrient concentration (Penttinen et al., 2018). In addition, putative virulence genes that increased in expression in rhizoid morphotype were different than the genes that were activated in rough morphotype as a response to increased resource level (Penttinen et al., 2016), highlighting the importance of inducible responses in pathogenesis in *F. columnare*.

In conclusion, our study demonstrated that fish farming environment induces phenotypic variation in growth in *F. columnare* as compared to natural waters and might therefore contribute to evolutionary trajectories in this bacterium. The high growth rate of the rough morphotypes of fish farm isolates especially in higher resource concentrations and in the higher temperature might be an adaptation to utilize increased resource concentrations accumulating in water at farms from uneaten fish feed and excreta due to impaired fish feeding in increased temperatures (Wedemeyer, 1996; Ellis et al., 2002). We did not find evidence for phenotypic variation in virulence based on the origin of the isolate. However, our data suggests that also the less virulent rough morphotype can quickly convert into virulent rhizoid type in contact with the fish. Combined with the higher growth rate of the rough morphotype at fish farms, morphotype reversibility is likely to increase the probability of columnaris epidemics at fish farms. Our results contribute to the growing evidence of the crucial role of intensive aquaculture in driving evolution in microbial pathogens (Mennerat et al., 2010; Pulkkinen et al., 2010; Sundberg et al., 2016; Mennerat et al., 2017).

## Experimental procedures

### The study species

*Flavobacterium columnare* exhibits three different colony morphologies, of which the rhizoid type (Rz) is associated with high virulence (Kunttu et al., 2009a). The other two types, rough (R) and soft (S) are less virulent (Kunttu et al., 2009a; Laanto et al., 2012). Primary isolations from natural samples on selective agar plates (Decostere et al., 1997) produce generally a rhizoid colony morphology, while the two other types appear during continued re-cultivation (Kunttu et al., 2009a; Sundberg et al., 2014), or as a response to infection by phages (Laanto et al., 2012).

### Isolation and cultivation of bacteria

Fish farm isolates were obtained from diseased fish or from tank water during outbreaks at fish farms as part of disease surveillance. The isolates from nature were collected from water or from a diseased wild fish upstream of a fish farm (Table 5). Bacteria isolations were performed with standard culture methods, using modified Shieh medium (from now on: Shieh medium) (Song et al., 1988) supplemented with tobramycin (Decostere et al., 1997). Pure cultures were stored at −80°C with 10% glycerol and 10% fetal calf serum.

**Table 5.**
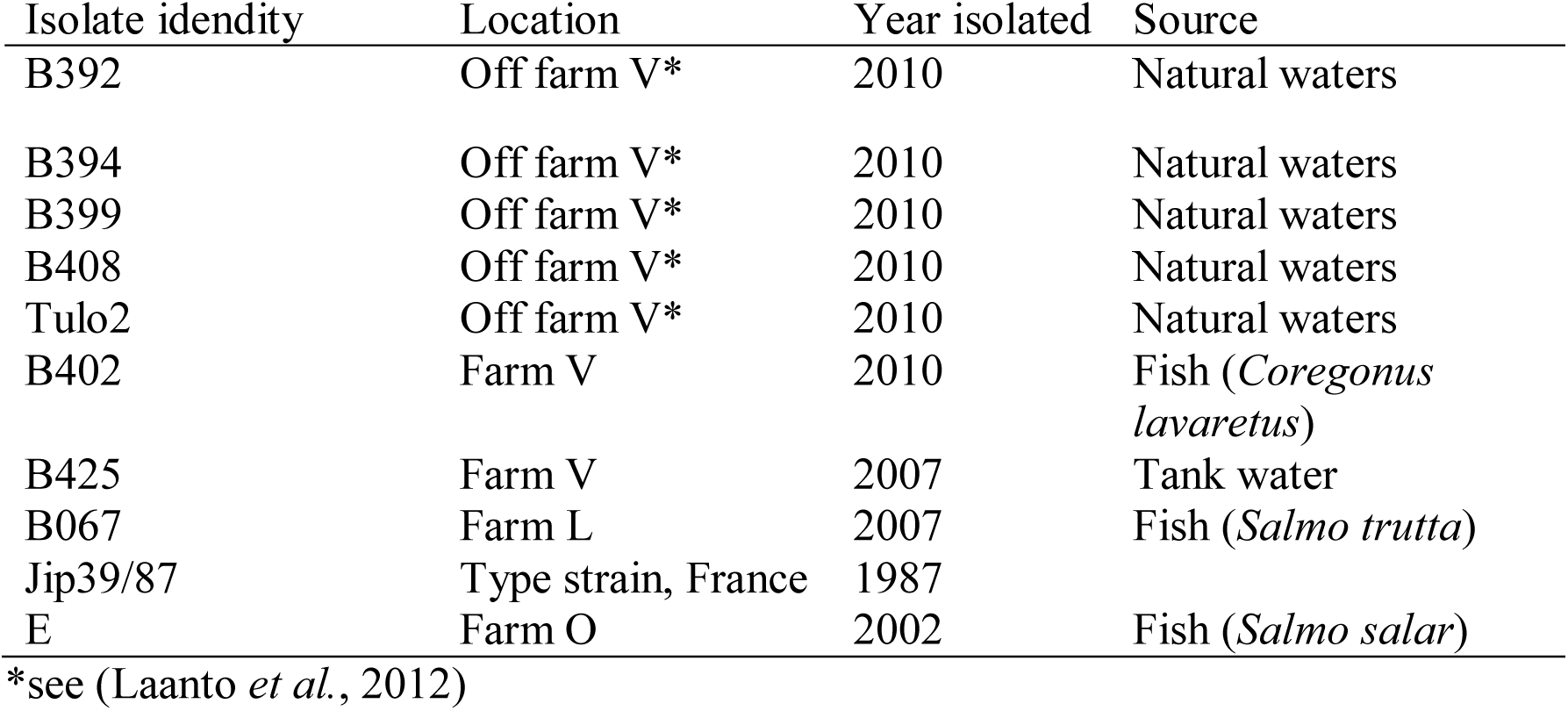
The *Flavobacterium columnare* isolates used in this study.

All isolates exhibited originally a rhizoid morphotype. Rough colonies that appeared spontaneously during plate cultivation were collected singly and cultured in Shieh medium at room temperature with constant shaking (210 rpm), and plated to assure the loss of parental Rz growth prior storing at −80°C (Fig. S1).

Prior to the experiments, bacteria were revived from frozen stocks by inoculation to Shieh medium (1x concentration). Bacteria were cultured at room temperature with constant agitation (210 rpm) for 24 h, and subsequently enriched in 1/10 dilution in fresh Shieh medium overnight in the same conditions. For growth measurements, 10 µL of the culture was applied into 100-well Bioscreen C^®^ plates containing 400 µL of fresh Shieh medium. Maximum growth rates and yield were determined from biomass growth data recorded at 5 min intervals with 420-580 nm optical density for 7 d at 25°C and 5 d at 15 °C. Measurements were conducted in five different resource concentrations (2x, 1x, 0.5x, 0.1x and 0.05x of Shieh medium). Each isolate-morphotype combination was included twice into separate measurements on two subsequent weeks, totaling to four replicate measurements. In addition, after liquid culture, bacterial samples were plate cultured on Shieh-agar with the five different resource concentrations mentioned above, to check the morphotype stability under different resource levels.

### Virulence experiments

The fish experiments were conducted according to the Finnish Act of the Use of Animals for Experimental Purposes, under permission ESAVI-2010-05569/Ym-23 granted by the National Animal Experiment Board at the Regional State Administrative Agency for Southern Finland for L-RS.

For testing virulence we used zebra fish, which have been established as a reliable model system for revealing differences in virulence among strains in *F. columnare* (Kinnula et al., 2015). Zebra fish (*Danio rerio*) were obtained from Core Facilities (COFA) and research services of Tampere University, Finland. The fish used in the experiment were disease-free, adult and unsexed. The size range of fish challenged with fish farm isolates and natural isolates were 0.06-0.88 g and 0.08-0.87 g, respectively. For each isolate-morphotype combination, 10 individual fish were challenged, with 10 additional fish used as controls, totaling in 210 individual fish. The fish were infected individually in continuous challenge (Kinnula et al., 2015) in 500 ml water with 1 × 10^5^ CFU ml^−1^ freshly grown bacteria added with 4 ml of fresh Shieh medium. For control fish, 4 ml of Shieh was added without bacteria. Fish were monitored for 4 d for disease symptoms and morbidity. Fish expressing disease symptoms were removed, put down by cutting the cordial spine with scissors and weighed. The presence of *F. columnare* infection on fish was checked with plating a primary culture from the gills onto modified Shieh agar plates.

### Data analysis

The maximum growth rates were assessed from the OD data by assuming that the population is growing at an exponential rate before the resources deplete. This assumption allows the estimation of the initial population size that is needed for calculating the population growth rate. The algorithm first smoothed the random fluctuations in the OD measurement values with 25 time point moving window and then used the log OD values for calculating the maximum rate of increase. The initial population size parameter that fulfilled best the assumption of exponential growth was used in the time series analysis. The 25°C data used all the data points as there were no apparent lag phases, whereas with the 15°C data the estimation of the maximum growth rate was started after 2 hours to exclude the lag phase.

As our dataset contained several factors that could interact in many ways, we utilized model selection criteria in order to reduce the complexity of the models explaining growth rate or yield. We started model building from model containing all possible interactions between measurement temperature (T), resource concentrations (R), origin of the isolate (O) and morphotype (M) (all fixed factors). In all models we included “maternal” clone identity as a random factor to control for non-independence resulting from four measurement replicates, and from the fact that colony types originated from the same bacterial culture. Moreover, the effect of measurement day was fitted as a fixed factor in all models. Starting from the highest order interactions, each of the interaction was removed from the model, if the interaction was not statistically significant (based on p-value<0.10) and its removal improved the model fit (AIC). Similar model selection was used to test how experimental factors explain biomass yield. Since yield is often found to trade off with growth rate (Velicer and Lenski, 1999) we tested also if our results concerning growth rate could be explained by inclusion of yield as a covariate. To explore if covariate has similar effects in all treatment groups we fitted also interactions of covariate (standardized to mean of zero) with all factors (Hendrix et al., 1982). Similar model selection as above was utilized to drop out non-significant factor to covariate interactions. Since covariate and factor by covariate interactions did not affect significance of our results or conclusions it is clear that changes between treatments are not caused by life-history tradeoffs between maximal growth rate and yield. Thus, these results are not presented or considered further. Statistical testing was performed with REML mixed models with IBM SPSS v. 19 (IBM).

In fish challenge experiments, the fish were monitored for 96 h and the last moribund fish was encountered and euthanized at 54 h. Thus the remaining challenged fish were considered as true survivors, and not as censored cases. Therefore, we used generalized linear mixed models for binomial distribution to examine the morbidity caused by different morphotypes of replicate strains originating from nature or from fish farm. We analyzed the morbidity risk of a host in an hour with a model including all possible interactions between origin of the isolate (O; nature, fish farm), colony type (T; rhizoid, rough) and fish weight (W; as a continuous covariate). Strain identity (S) was included as a random factor. Model reduction based on AIC criteria was performed with drop1-function in package MASS (Venables and Ripley, 2002) starting from the full model. The analysis was conducted using R software (version 3.3.2) and Lme4 package (R Development Core Team, 2015).

## Acknowledgements

The authors want to thank J.-F. Bernardet for providing Fc isolate JIP39/87 and H. Kunttu for isolates from natural waters used in the study and M. Nicolini, H. Kinnula and R. Penttinen for assistance in the laboratory. This study was supported by The Centre of Excellence in Biological Interactions (research themes led by Prof. Jaana K. Bamford and Prof. Johanna Mappes, #252411) and by Finnish Cultural Foundation (KP), by Academy of Finland (grants #266879, #304615, #7128888, #314939), and by Jane and Aatos Erkko foundation.

