## Supplementary material for "Rich resource environment of fish farms facilitates phenotypic variation and virulence in an opportunistic fish pathogen": Figure S1

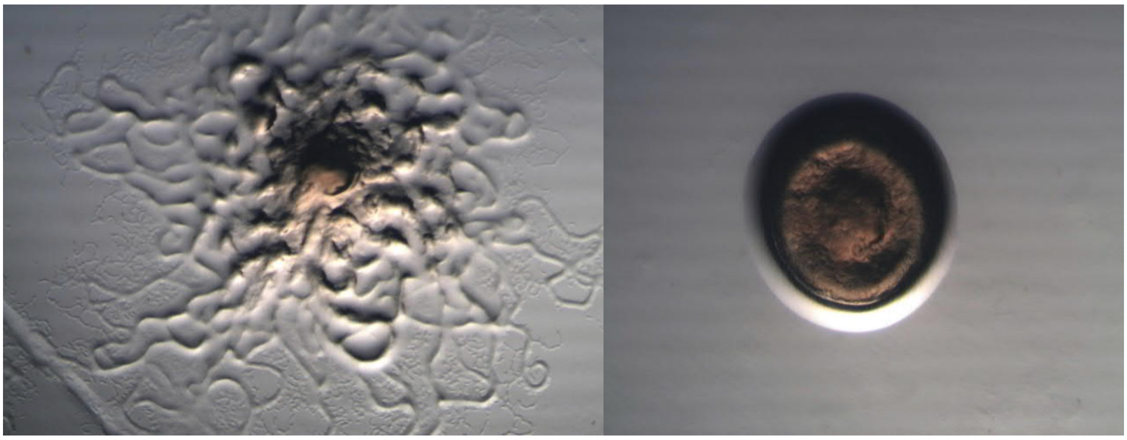

Figure S1. A rhizoid (left) and a rough (right) morphotype of *Flavobacterium columnare* cultured on Shieh-agar plates.

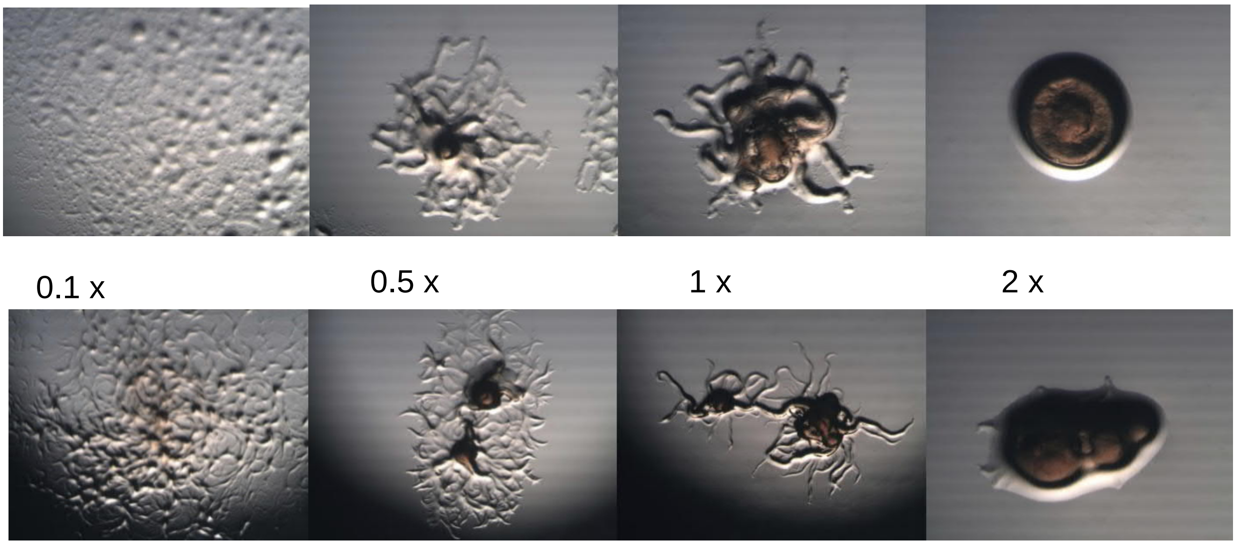


Figure S2. Examples of colony morphologies of *Flavobacterium columnare* plated on agar plates containing different concentrations of Shieh medium (0.1 x, 0.5  x, 1 x and 2x Shieh). Upper panel isolate B067, lower panel isolate E.
